# A high-quality sequence of *Rosa chinensis* to elucidate genome structure and ornamental traits

**DOI:** 10.1101/254102

**Authors:** L. Hibrand Saint-Oyant, T. Ruttink, L. Hamama, I. Kirov, D. Lakwani, N.-N. Zhou, P.M. Bourke, N. Daccord, L. Leus, D. Schulz, H. Van de Geest, T. Hesselink, K. Van Laere, S. Balzergue, T. Thouroude, A. Chastellier, J. Jeauffre, L. Voisine, S. Gaillard, T.J.A. Borm, P. Arens, R.E. Voorrips, C. Maliepaard, E. Neu, M. Linde, M.C. Le Paslier, A. Bérard, R. Bounon, J. Clotault, N. Choisne, H. Quesneville, K. Kawamura, S. Aubourg, S. Sakr, M.J.M. Smulders, E. Schijlen, E. Bucher, T. Debener, J. De Riek, F. Foucher

## Abstract

Rose is the world’s most important ornamental plant with economic, cultural and symbolic value. Roses are cultivated worldwide and sold as garden roses, cut flowers and potted plants. Rose has a complex genome with high heterozygosity and various ploidy levels. Our objectives were (*i*) to develop the first high-quality reference genome sequence for the genus *Rosa* by sequencing a doubled haploid, combining long and short read sequencing, and anchoring to a high-density genetic map and (*ii*) to study the genome structure and the genetic basis of major ornamental traits.

We produced a haploid rose line from *R. chinensis* ‘Old Blush’ and generated the first rose genome sequence at the pseudo-molecule scale (512 Mbp with N50 of 3.4 Mb and L75 of 97). The sequence was validated using high-density diploid and tetraploid genetic maps. We delineated hallmark chromosomal features including the pericentromeric regions through annotation of TE families and positioned centromeric repeats using FISH. Genetic diversity was analysed by resequencing eight *Rosa* species. Combining genetic and genomic approaches, we identified potential genetic regulators of key ornamental traits, including prickle density and number of flower petals. A rose *APETALA2* homologue is proposed to be the major regulator of petals number in rose. This reference sequence is an important resource for studying polyploidisation, meiosis and developmental processes as we demonstrated for flower and prickle development. This reference sequence will also accelerate breeding through the development of molecular markers linked to traits, the identification of the genes underlying them and the exploitation of synteny across *Rosaceae*.

## Background

Rose is the queen of flowers, and holds great symbolic and cultural value. Roses appeared as decoration on 5000 year-old Asian pottery [1], and Romans cultivated roses for their flowers and essential oil [2]. Today, there are no ornamental plants with greater economic importance than roses. They are cultivated worldwide and sold as garden plants, in pots, or as cut flowers, the latter accounting for approximately 30% of the market. Roses are also used for scent production and for culinary purposes [3].

Roses present an ideal model for woody and ornamental plants, but also display a range of unique features thanks to their complex evolutionary history including interspecific hybridization events and polyploidisation [4–6]. Roses belong to the genus *Rosa (Rosoideae, Rosaceae)*, which contains more than 150 species [7] with varying levels of ploidy, ranging from 2n=2× to 10× [8, 9]. Many modern roses are tetraploid and segregate as ‘segmental’ allopolyploids; a mixture between allopolyploidy and autopolyploidy [10], while dog-roses display unequal meiosis to maintain pentaploidy [11]. Selection and breeding of roses has a long, yet mostly unresolved, history in Europe and Asia, which most likely involved several interspecific hybridization events. Due to the strong and continuous interest, even very old varieties have been conserved in private and public rose gardens, and represent a living historical archive of the breeding and selection process [12]. In addition, large and well-documented herbarium collections, combined with the advent of advanced genomic analysis, offer excellent opportunities to reconstruct the underlying phylogenetic relationships.

Performance-related traits selected for in roses are different from agronomic traits in field crops. Production and resistance to biotic and abiotic stresses are important, but aesthetic criteria have played an essential role during the last 250 years of rose selection and breeding, including colour of the flower, architecture of the flower ranging from simple flowers with five petals to ‘double’ flowers with over 100 petals, biosynthesis and emission of volatile molecules producing the typical scent, and formation of prickles on the stem and leaves. Although important for domestication, ornamental traits primarily serve adaptation to natural conditions. The availability of a high quality reference genome sequence is key to unravel the genetic basis underlying these evolutionary and developmental processes, as it will accelerate future genetic, genomic, transcriptomic and epigenetic analyses. Recently, a draft reference genome sequence of *Rosa multiflora* has been published [13]. Whereas completeness measures suggest that the assembly is fairly complete in terms of the gene space covered, it is also highly fragmented (83189 scaffolds, N50 of 90kbp).

Here, we present an annotated high-quality reference genome sequence for the *Rosa* genus using a haploid rose line derived from an old Chinese *Rosa chinensis* variety ‘Old Blush’ (Figure 1a). ‘Old Blush’ (syn. Parsons’ Pink China), which was introduced to Europe and North America in the 18th century from China, is one of the most influential genotypes in the history of rose breeding. ‘Old Blush’ was important for introducing recurrent flowering, an essential trait for the development of modern rose cultivars [14]. Our pseudo-chromosome scale genome assembly was validated using both high-density genetic maps of multiple F1 progenies and synteny with *Fragaria vesca*. We delineated hallmark chromosomal features such as the pericentromeric regions through annotation of TE families and positioning of centromeric repeats using FISH. This reference genome allowed a detailed analysis of genetic diversity within the *Rosa* genus following a resequencing of eight wild species. Using this reference genome sequence combined with genetic (F1 progeny and diversity panel) and genomic approaches, we were able to identify key potential genetic regulators of important ornamental traits: continuous flowering, flower development, prickle density and self-incompatibility.

**Figure 1.**
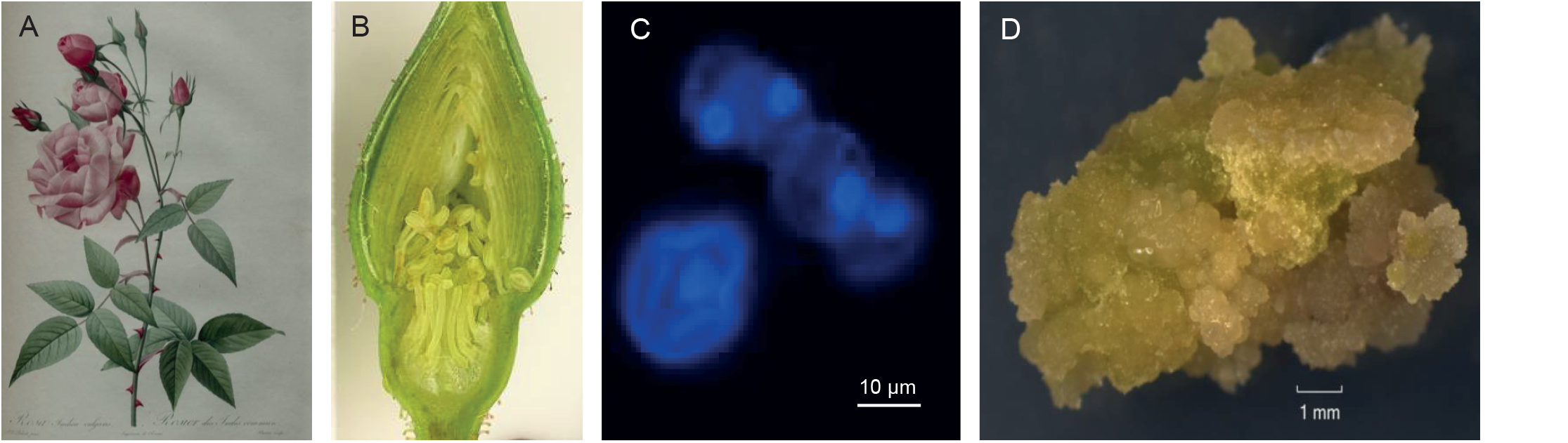
Development of the HapOB haploid line from *R. chinensis* ‘Old Blush’. A) *R. chinensis* variety ‘Old Blush’ painted by Redouté in 1814. B) Cross section of floral stage used for anther culture. C) DAPI staining on midto late uninucleate microspores. D) HapOB callus obtained after microspore culture, used for genome sequencing.

## Results

### (1) Development of a high-quality reference genome sequence

We have developed a haploid callus cell line (HapOB) using anther culture at mid-to late uninucleate microspores developmental stage from the diploid heterozygous ‘Old Blush’ variety (Figure 1b, 1c and 1d). The homozygosity of the HapOB line was verified with 10 microsatellite markers distributed over the seven linkage groups (Supplementary Table 1). Flow cytometric analysis showed that the HapOB callus is diploid, suggesting that spontaneous genome doubling occurred during *in vitro* propagation.

A combination of Illumina short-read sequencing and PacBio long-read sequencing technologies was used to assemble the doubled-haploid HapOB genome sequence. PacBio sequencing data (1.64 M reads with an N50 of 20.0 kbp; total 19.3 Gbp; 40× genome coverage), was assembled with CANU [15], yielding 551 contigs (N50 of 3.4 Mbp, L75 of 97) representing a total length of 512 Mbp. 95% of the obtained sequence is contained in only 196 contigs. The PacBio-based assembly was error-corrected with Illumina paired-end reads (37.3k SNPs and 307.7k indels were corrected, representing 341.1 kbp). K-mer spectrum analysis (K=25) suggested a genome size of 532.7 Mbp (251.1 Mbp of unique genome sequence and 279.6 Mbp of repetitive sequences), while flow cytometric analysis estimated a genome size of 1C=568±9 Mbp. The assembled sequence therefore represents 96.1% or 90.1%, respectively, of the estimated genome size.

High density female and male genetic maps were developed from a cross between *R. chinensis* ‘Old Blush’ and a hybrid of *R. wichurana* (OW) of which 151 F1 progeny were genotyped with the 68K WagRhSNP Axiom array ([16], Table 1 and Supplementary Table 2). Thirteen contigs where marker order clearly indicated assembly artefacts were split before anchoring all 564 resulting contigs to the female and male genetic maps using a total of 6746 SNP markers (Table 1). Of these, 196 contigs were anchored manually onto the seven linkage groups (LG), mostly on both the female and male genetic maps (174 and 143 contigs, respectively). In total, 466 Mbp were thus anchored onto the genetic maps and assembled into seven pseudo-chromosomes representing 90% of the assembled contig length (Table 1 and Supplementary Figure 1a). The remaining 368 contigs (52 Mbp) were assigned to Chr0. The quality of the assembly of the seven pseudo-chromosomes was assessed using two independent genetic maps: the previously published integrated high-density genetic map (K5) based on 25695 SNPs in tetraploid rose [10], and a newly developed diploid genetic map based on 174 F1 progeny from a cross between cultivar ‘Yesterday’ and *R. wichurana* (YW, see Supplementary Figure 1b). The co-linearity between the pseudo-chromosomes and the respective linkage maps is excellent (Supplementary Figure 2). In addition, anchoring of the 386 contigs (52 Mbp) currently assigned to Chr0, onto the tetraploid K5 map and the YW map revealed that, respectively, 39 contigs (total 28.4 Mbp) and 27 contigs (total 24.1 Mbp), can potentially be positioned onto the seven linkage groups (Supplementary Figure 2). However, because these genetic maps were created using independent genotypes not related to *R. chinensis* ‘Old Blush’, we took a conservative approach and did not yet incorporate these contigs into the pseudo-chromosome sequence of the HapOB genotype.

**Table 1:** Metrics of the alignment of the male and female genetic maps with the HapOB genome assembly. The genetic maps were developed from a cross between ‘Old Blush’ (female) and a hybrid of *R. wichurana* (male) using a Affymetrix SNP array. The initial size of the genome was 512Mb, and reached finally 518,5 due to the addition of 10000N between each contigs to create the pseudomolecules.

| LG | Genetic maps (No. of markers) |  | Chr. | No. of anchored markers used for anchoring |  | No. of anchored contigs |  |  |  |  | Pseudo-molecules |
| --- | --- | --- | --- | --- | --- | --- | --- | --- | --- | --- | --- |
|  | female (OB) | male (W) |  | female | male | female | male | manual integration | Cut | Excluded | Size (in bp) |
| 1 | 715 | 195 | 1 | 587 | 146 | 18 | 14 | 18 | 1 | 1 | 64 770 848 |
| 2 | 1114 | 303 | 2 | 1001 | 249 | 14 | 18 | 20 |  |  | 75 129 302 |
| 3 | 528 | 564 | 3 | 477 | 498 | 20 | 25 | 31 |  | 1 | 46 843 630 |
| 4 | 227 | 404 | 4 | 191 | 334 | 12 | 18 | 20 |  |  | 59 004 735 |
| 5 | 1031 | 362 | 5 | 866 | 275 | 40 | 29 | 37 | 2 | 1 | 85 885 663 |
| 6 | 1153 | 254 | 6 | 1010 | 186 | 43 | 20 | 43 |  | 1 | 67 395 200 |
| 7 | 863 | 241 | 7 | 743 | 183 | 27 | 19 | 27 |  |  | 67 081 725 |
|  |  |  |  | **Total without Chr 0** |  | **174** | **143** | **196** |  |  | **466 111 103** |
|  |  |  | 0 | - | - | 387 | 418 | 368 |  |  | 52 404 850 |
| **Total:** | **5631** | **2323** | **Total** | **4875** | **1871** | **561** | **561** | **564** |  |  | **518 515 953** |

### (2) Positioning centromeres within the genome assembly

Bioinformatics and cytogenetics were used to identify the centromeric regions. We discovered a highly abundant tandem repeat (0.06% of the genome with more than 2000 copies per haploid genome) with a 159 bp monomer length that we call OBC226 (‘Old Blush’ centromeric repeat from RepeatExplorer cluster 226, Figure 2a). PCR confirmed the tandem organization of this repeat (Figure 2b). FISH analysis unambiguously confirmed the location of the repeat in the centromeric regions of four out of seven chromosomes, i.e. Chr2, Chr5, Chr6 and Chr7 (Figure 2c). Mapping of the OBC226 repeat sequence revealed regions with high coverage on all HapOB pseudo-chromosomes except Chr1, explaining why no clear centromeric region could be detected on this Chromosome (Figure 2d). On the other two chromosomes, Chr3 and Chr4, the copy number of OBC226 was likely too low to be detected by FISH. Furthermore, the core OBC226 centromeric repeats were flanked by other repetitive sequences, and these were unequally distributed along the chromosomes, with a clearly higher density in the core centromeric regions (Figure 2d). These centromeric regions were also enriched in Ty3/Gypsy transposable elements. Taken together, these results confirm the position of the centromeric regions in the seven pseudo-chromosomes and reveal the high repeat sequence content, and low gene content, of the scaffolds currently assigned to Chr0.

**Figure 2.**
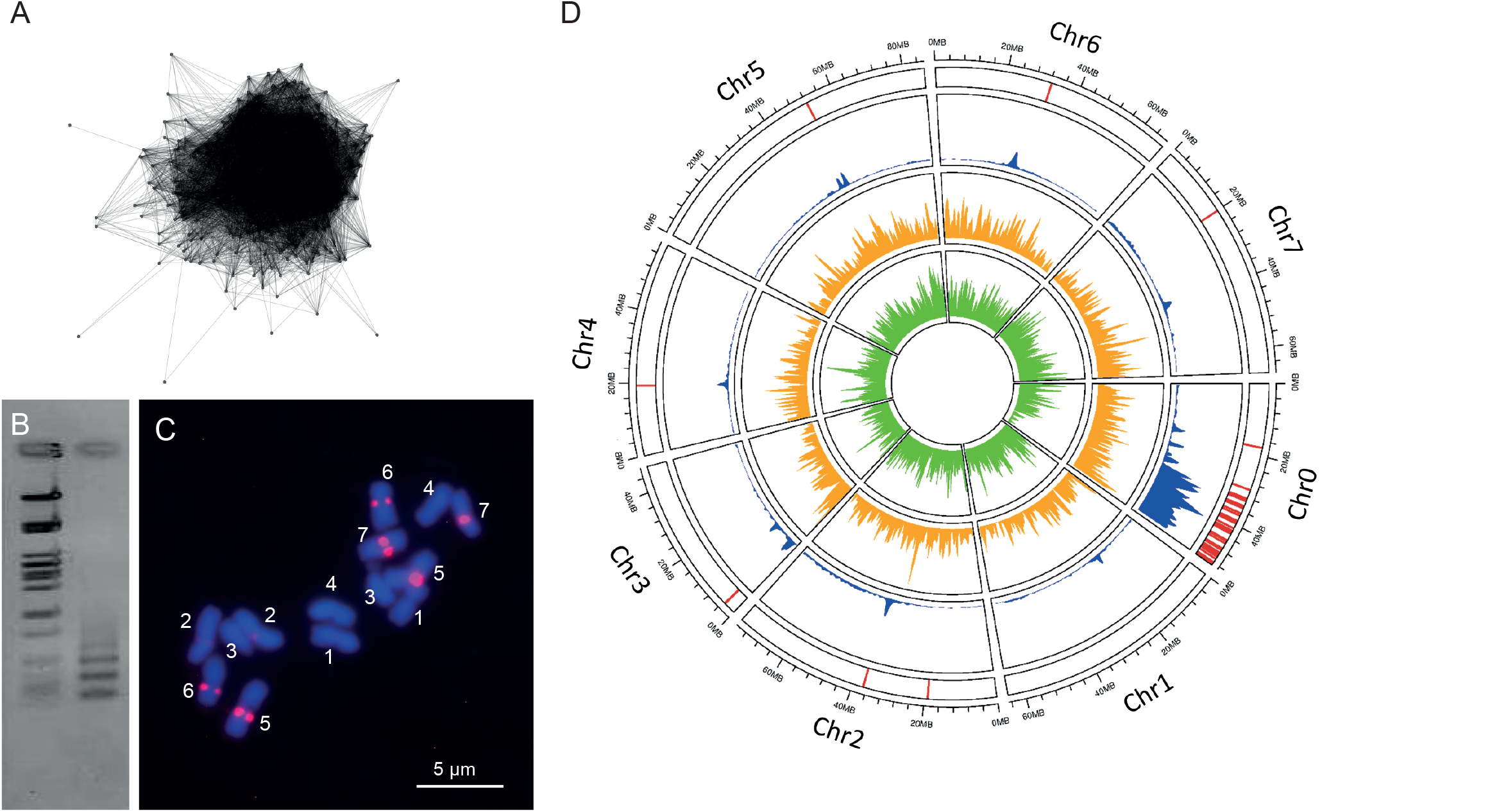
Identification of centromeric regions in the HapOB reference genome. A) cluster CL226 identified by RepeatExplorer. B) Agarose gel-electrophoresis of tandem repeat fragments amplified from genomic DNA of HapOB using OBC226 PCR-primers (right lane) along with lambda-*Pst*I size ladder (left lane). C) FISH with TAMRA-labeled OBC226 oligo probes on *R. chinensis* metaphase chromosomes. Chromosome numbers are labeled as 1-7. D) Circos representation of the distribution of OBC226 (red), pericentromeric region (blue), Ty3/Gypsy (yellow), and Ty1/Copia repeat elements (green) along the seven pseudo-chromosomes and Chr0.

### (3) Annotation of the sequence

#### Coding genes

Based on the mapping of 723,268 transcript sequences (EST/cDNA and RNA-seq contigs with a minimum size of 150 bp) onto the HapOB genome assembly, we predicted a total of 44,481 genes covering 21% of the genomic sequence length using Eugene combiner [17]. These include 39,669 protein coding genes and 4,812 non-coding genes. Evidence of transcription was found for 87.8% of all predicted genes. At least one InterPro domain signature was detected in 86.5% of the protein-coding genes using InterProScan [18] with 68.0% of the genes assigned to 4,051 PFAM gene families [19]. The quality of the structural annotation was assessed using the BUSCO v2 method based on a benchmark of 1440 conserved plant genes [20], of which 92.5% had complete gene coverage, 4.1% were fragmented and only 3.4% were missing. The set of predicted non-coding genes included 186 rRNA, 751 tRNA, 384 snoRNA, 99 miRNA, 170 snRNA and 3,222 unclassified genes (annotated as ncRNA) with transcription evidence but no consistent coding sequence.

#### Transposable elements

The REPET package [21] was used to produce a genome-wide annotation of repetitive sequences of the HapOB genome (see Materials and Methods for details). Retrotransposons, also called class I elements, cover the largest TE genomic fraction (35,1% of the sequenced genome), with LTR-retrotransposons representing 28.3% and Gypsy covering a larger part than Copia (Supplementary Table 3 and Supplementary Figure 3). Non-LTR retrotransposons (LINE and potential SINE) represent 5,0 % and class II elements (DNA transposons and Helitrons) 11.7% (Supplementary Figure 3). The remaining 15.1% include unclassified repeats (7.3%), chimeric consensus sequences (1.9%) and potential repeated host genes (5.8%) kept in this study. We also identified caulimoviridae copies representing 1.25% of the genome. Interestingly, one particularly-abundant Gypsy Tat-like family was found in the genome assembly. The total copy coverage represents 3.4% of the genome. Tat-like elements are known to have an ORF after the polymerase domains and surprisingly in this case the ORF corresponds to a class II transposase domain.

### (4) Synteny between *Rosa* and *Fragaria vesca*

*Rosa* and *Fragaria* are closely related as they both belong to the *Rosoideae* subfamily of the *Rosaceae* [22]. Previous genetic studies have demonstrated that large macro-syntenic blocks are conserved between *Rosa* and *Fragaria* [10, 23]. We compared the HapOB genome to the previously published *Fragaria* genome [24] to analyse synteny in more detail (Supplementary Figure 4). At the macro-synteny level, large blocks are conserved between *Rosa* chromosomes 4, 5, 6 and 7 and *Fragaria* chromosomes 4, 3, 2 and 5, respectively. *Rosa* Chromosome 1 is largely syntenic to *Fragaria* Chromosome 7. Consistent with previous suggestions [10], a reciprocal translocation was detected between *Rosa* chromosomes 2 and 3 and *Fragaria* chromosomes 6 and 1, respectively, but synteny can also be detected between *Fragaria* Chromosome 1 and *Rosa* Chromosome 6 (Supplementary Figure 4). However, the composition of *Fragaria* Chromosome 1 looks more complex than previously thought, with conserved blocks from *Rosa* chromosomes 2, 3, 6 and 7, and some segments of *Rosa* chromosomes 3 and 7 are syntenic with regions of *Fragaria* Chromosome 7. Our results provide a more complete picture of the synteny between rose and strawberry than previous attempts [10, 23], highlighting numerous small rearrangements as proposed between strawberry, peach and apple [25].

### (5) Genetic diversity with the genus *Rosa*

The more than 150 existing rose species belong to four subgenera. Except for the subgenus *Rosa*, the other three only contain one or two species. We resequenced eight *Rosa* species, representing three subgenera (*Hulthemia: R. persica, Herperhodos: R. minutifolia, Rosa: R. chinensis* var. *spontanea, R. rugosa, R. laevigata, R. moschata, R. xanthina spontanea* and *R. gallica*). The six genotypes of subgenus *Rosa* represent different sections according to the latest phylogenetic analyses (Table 2) [26, 27]. We identified SNPs and InDels relative to the HapOB reference sequence (Figure 3). The lowest SNP and Indel density was found to be in *R. chinensis* var. *spontanea* (9.9 and 1.6 per kbp respectively), whereas the highest was found to be in *R. gallica* (21.0 and 4.5 per kbp respectively). As expected, the majority of SNPs (range 79.2-89.0%) are located in non-coding sequences (downstream, upstream and intergenic regions, Supplementary Table 4). 3-7% of the SNPs are located in exons, of which half have a moderate (synonymous) or high effect on the protein sequence, in line with previous observations in other species (as tomato [28]). Furthermore, the different species display varying levels of homozygosity with ratios of heterozygous to homozygous SNPs ranging from 79.2% in *R. persica* to 26.0% in *R gallica* (Supplementary Table 4).

**Table 2:** Summary of resequencing and variations (SNPs and Small Indels) indentifed in 8 *Rosc*

| *Rosa* species sequenced | Genome Size<br>(in Mbp) | Classification |  | ploid<br>y | No. of reads | No. of reads<br>mapped | No. SNPs | SNP /<br>density<br>(no./kbp) | No. Small<br>Indels | Small Indel<br>density<br>(no./kbp) |
| --- | --- | --- | --- | --- | --- | --- | --- | --- | --- | --- |
|  |  | Subgenus | Section |  |  |  |  |  |  |  |
| *R. chinensis* var *spontanea* | 562 | *Rosa* | *Chinenses* | 2 | 110 470 873 | 104 074 609 | 5 564 345 | 9,9 | 876 648 | 1,6 |
| *R. gallica* | 538 | *Rosa* | *Gallicanae* | 4 | 230 772 574 | 218 080 082 | 11 280 831 | 21,0 | 2 430 138 | 4,5 |
| *R. laevigata* | 562 | *Rosa* | *Laevigatae* | 2 | 100 084 132 | 92 357 637 | 6 327 292 | 11,3 | 1 195 164 | 2,1 |
| *R. moschata* | 554 | *Rosa* | *Synstylae* | 2 | 92 392 169 | 86 118 741 | 5 862 043 | 10,6 | 1 417 766 | 2,6 |
| *R. munitifolia* *alba* | 416 | *Hesperodos* |  | 2 | 95 941 308 | 89 100 693 | 5 270 249 | 12,7 | 1 208 933 | 2,9 |
| *R. persica* | 416 | *Hulthemia* |  | 2 | 113 791 888 | 100 375 824 | 5 602 086 | 13,5 | 1 218 337 | 2,9 |
| *R. rugosa* | 522 | *Rosa* | *Rosa* | 2 | 125 451 864 | 115 867 342 | 8 270 874 | 15,8 | 1 703 127 | 3,3 |
| *R. xanthina* *spontanea* | 391 | *Rosa* | *Pimpinellifolia* | 2 | 94 747 185 | 84 959 801 | 5 642 595 | 14,4 | 1 316 384 | 3,4 |

**Figure 3.**
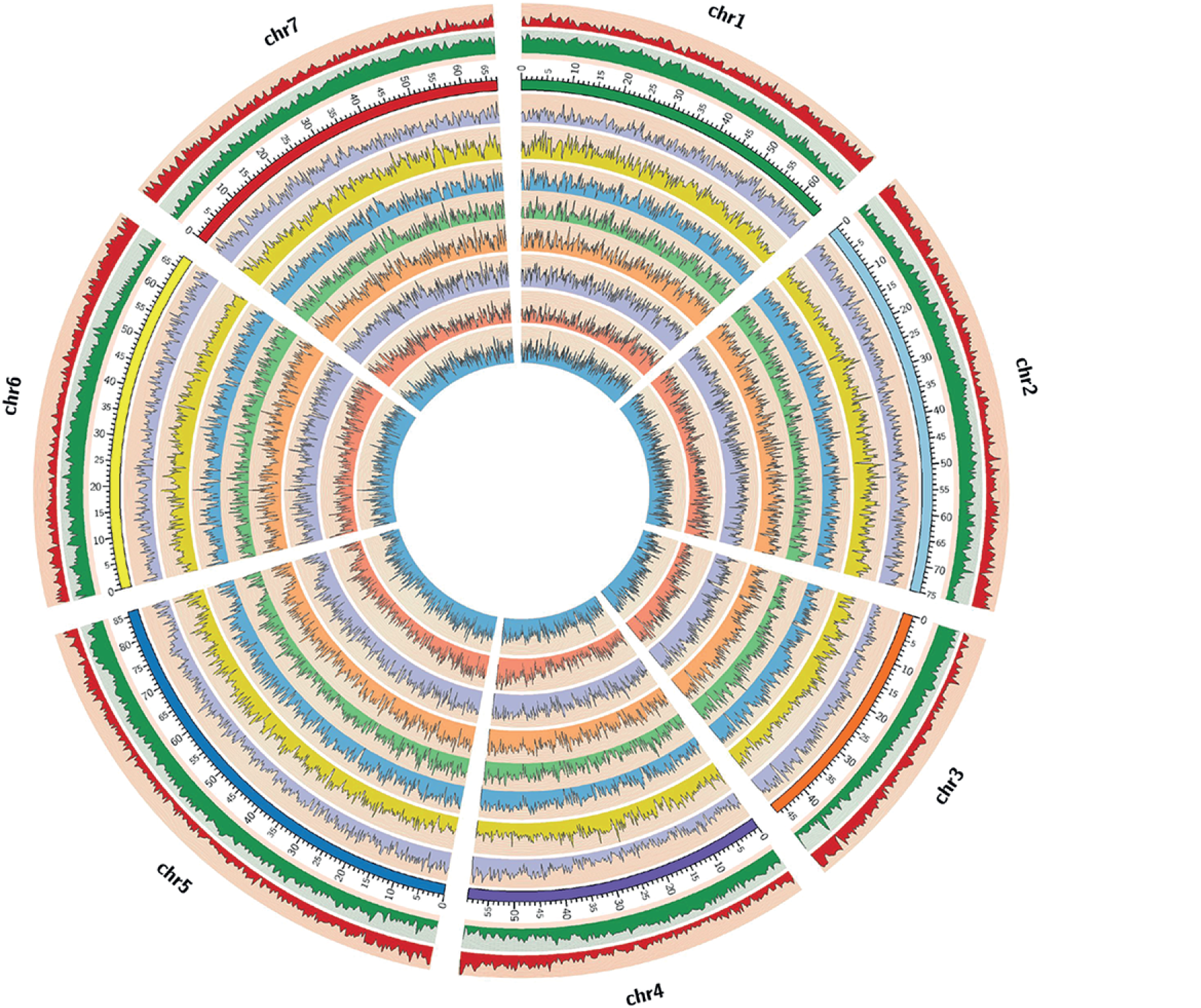
Analysis of genetic diversity in eight species of the *Rosa* genus along the seven pseudo-chromosomes of the HapOB reference sequence. Circles from outside to inside show: gene density (red), Transposable Element (TE) density (green), SNP density for *R. xantina* (blue), *R. chinensis* var. *spontanea* (yellow), *R. gallica* (blue), *R. laevigata* (green), *R. moschata* (orange), *R. rugosa* (blue), *R. persica* (pink), and *R. minutifolia* (blue). Scales are in Mbp.

Among these eight genomes, the closest relative to ‘Old Blush’ is *R. chinensis* var. *spontanea*. ‘Old Blush’ is described as an interspecific cross between *R. chinensis* var. *spontanea* and *R*. x *odorata* var *gigantea* [6], consistent with the relatively low sequence divergence of *R. chinensis* var. *spontanea* compared to the HapOB reference sequence. The high degree of sequence diversity in *R. gallica* could be due to tetraploidy, as shown by its high proportion of heterozygous SNP (74%, Supplementary Table 4). The number of small InDels was higher (between 876,648 and 2,430,123) compared to *Malus* with an average of 346,498 Indels [29], suggesting a higher level of diversity within the *Rosa* genus.

### (6) Analysis of the genetic determinism of important traits

This new reference sequence is an important tool to decipher the genetic basis of ornamental traits such as blooming (including continuous flowering, flower development and number of petals), prickle density on the stem and self-incompatibility. We studied the genetic determinism (*i*) in two F1 progenies (151 individuals, obtained from a cross between *R. chinensis* ‘Old Blush’ and a hybrid of *R. wichurana* (OW) and 174 individuals obtained from a cross between ‘Yesterday’ and *R. wichurana* (YW)) and (*ii*) in a panel of 96 rose cultivars originating from the 19^th^ to the 21^st^ century [30, 31]. Our data demonstrate that important loci controlling continuous flowering, double flower morphology, self-incompatibility and prickle density were predominantly localised on a genomic region on Chromosome 3 (Figure 4a).

**Figure 4.**
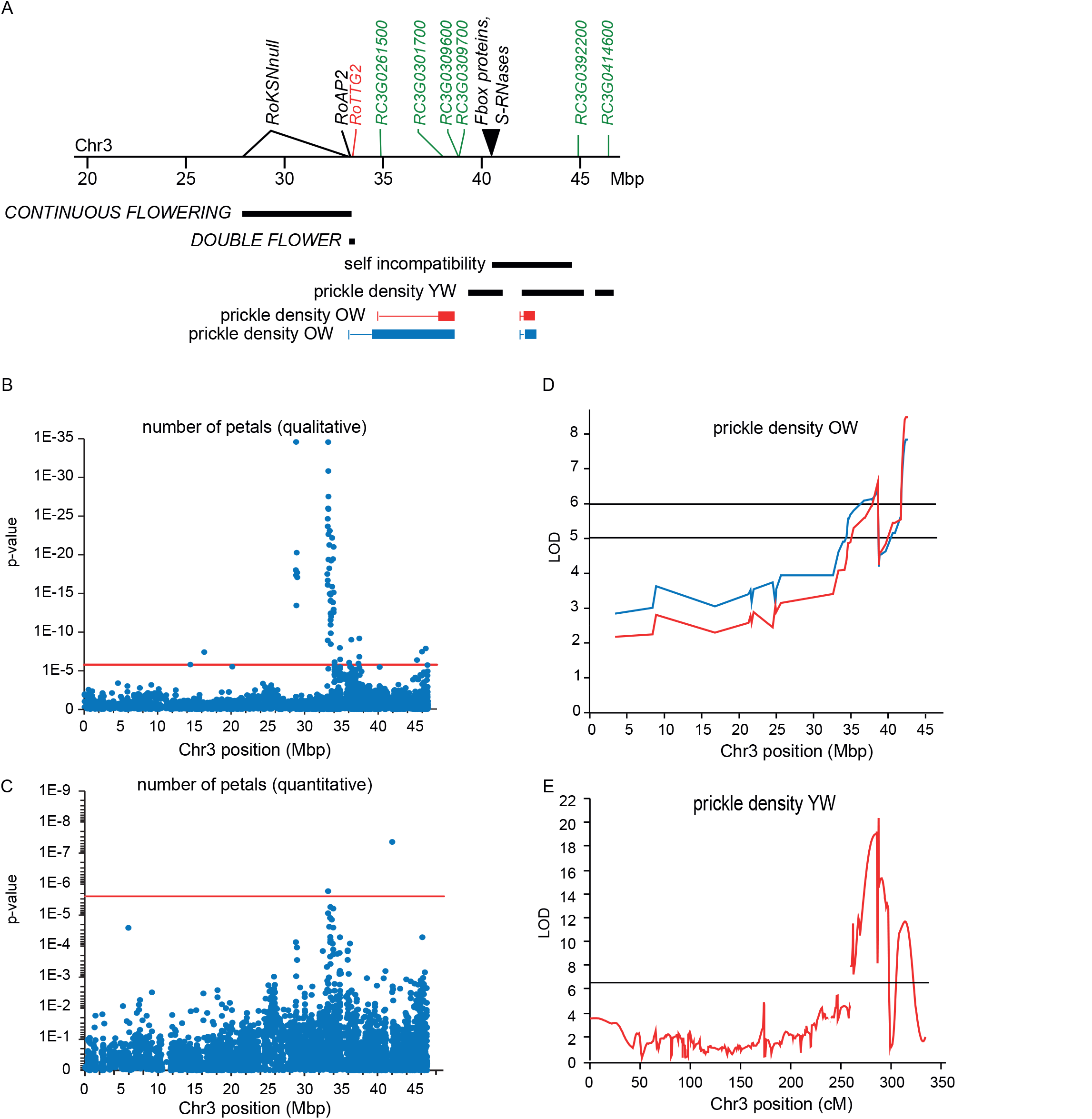
A region at the end of Chromosome 3 (Chr3) controls important ornamental traits. A) Major genes and QTLs that control continuous flowering, double flower, self-incompatibility and prickle density are shown together with candidate genes for each trait. Detailed analysis per locus are described in Supplemental Figures 5, 7, 9 and 10, respectively. B-C) GWAS analysis showing p-values of the association between SNPs positioned along Chr3 and the number of petals, indicating regions that control the number of petals; with petals considered as a B) qualitative trait (simple versus double) or C) quantitative trait. Red line represents Bonferroni corrected level of significance by number of contigs (for qualitative analysis, 19083 contigs = 2.48E-6; for quantitative analysis, 28054 contigs = 1.88E-6). D-E) QTL analysis for prickle density in two F1 progenies: D) the OW mapping population based on scoring from 2016 (blue line) and 2017 (red line); and E) the YW mapping population.

#### A. Detection of a new allele controlling continuous flowering in rose

Most species roses are once flowering (OF). In rose, continuous flowering (CF) is controlled by an homolog of the *TERMINAL FLOWER 1* (*TFL1*) family, *RoKSN*, which is located on LG3 [32]. The CF phenotype is due to the insertion of a *Copia* retrotransposon element in the *RoKSN* gene. The CF rose ‘Old Blush’ was previously proposed to be *RoKSN^copia^/RoKSN^copia^* at the *RoKSN* locus [32]. Here, using new QTL analyses on the OW progeny, we again identified the *CF* locus on LG3 (Figure 4a), but we were unable to detect the *RoKSN* gene in the annotated HapOB genome. Detailed analysis of *RoKSN* allele segregation in the OW progeny revealed the existence of a null allele, in which *RoKSN* is deleted (details in Supplementary Table 5). The diploid ‘Old Blush’ parent of the OB mapping population is therefore hemizygous *RoKSN^copia^/RoKSN^null^*, and the *RoKSN^null^* allele is present in the HapOB genome sequence.

Interesting parallels exist between rose and *F. vesca* because *F. vesca* also exhibits both OF and CF phenotypes. In strawberry a 2 bp deletion in the *TFL1* homolog causes a shift from OF to CF [32]. Synteny analysis revealed four orthologous syntenic blocks in the *RoKSN* gene region, here called block A-D (Supplementary Figure 5). We detected conserved gene content with genome rearrangements between different *Rosa* species and the published genome sequence of *Fragaria vesca* [24] where the synteny with *F. vesca* is broken at the *FvKSN* location. The *FvKSN* gene is located just between the A and B blocks in *F. vesca*. The A block is inverted in the HapOB genome, and the C and D blocks are inserted between the A and B blocks. In *Rosa multiflora* [13] and in *R. laevigata* (see Materials and Methods for the partial *R. laevigata* genome sequence assembly), which are both OF rose species, the *RoKSN^WT^* allele is present and synteny is conserved with *F. vesca* (Supplementary Figure 5). Taken together, these data suggest that the *RoKSN^null^* allele is the result of a large rearrangement at the *CF* locus leading to the complete deletion of the *RoKSN* gene. The *RoKSN^null^* allele represents a novel allele responsible for continuous flowering which has not been previously described.

#### B. Double flower

It was previously shown that the genetic basis of the double flower trait in rose is complex, with a dominant gene (*DOUBLE FLOWER*) controlling simple *versus* double flower phenotypes and two QTLs controlling the number of petals on double flowers [33]. Here, we combined the genome sequence with segregation data of four different F1 progenies to confine the putative location of the *DOUBLE FLOWER* locus (Supplementary Table 6) to a region of 293 kbp (between position 33.24 Mbp and 33.53 Mbp, Figure 4a). By GWAS analysis on 96 cultivated roses, we also detected a strong association between SNP and simple vs double flower GWAS analysis in the same region (between position 33.08 Mbp and 33.94 Mbp, Figure 4b); a second significant peak was located at a distance of 5 Mbp.

The 293 kbp region contains 41 annotated genes. Among these, half are expressed during the early stages of floral development in rose (Supplementary Table 7). Furthermore, by excluding genes that are expressed in late stages (complete open flower), we retained four genes: an F-box protein (RC3G0245100), a homologue of *APETALA2* (RC3G024300), a Ypt/Rab-GAP domain of gyp1p superfamily protein (RC3G0245000) and a tetratricopeptide repeat (TPR)-like superfamily protein (RC3G0243500) (Supplementary Table 7).

Concerning double flowering, ‘Old Blush’ is heterozygous for the *DOUBLE FLOWER* locus. The recessive *double flower* allele is coupled with the *RoKSN^null^* allele, suggesting that in HapOB the recessive allele has been sequenced. Sequencing both alleles of the four selected candidate genes in the original heterozygous diploid ‘Old Blush’ revealed only minor modifications for RC3G0245100, RC3G0245000 and RC3G0243500 (Supplementary Figure 6a, b and c respectively). Concerning the *APETALA2* gene (RC3G0243000), we were unable to completely amplify the second allele by PCR due to a large insertion in intron eight (Supplementary Figure 7a). The flanking sequences of this insertion showed similarity with an LTR gypsy retrotransposon (RLG_denovoHM-B-R10791_Map9) or an unclassified repeat element (noCAT_denovoHM-B-R7962) (Supplementary Table 3). No other differences were detected (except for a few SNPs) between the two alleles. Phylogenetic analysis showed that RC3G0243000 belongs to the *APETALA2* clade within the *AP2/ERF* subfamily [34] (Supplementary Figure 7d). Like all members of the AP2 clades, the protein encoded by RC3G0243000 contains two conserved AP2 domains and a conserved putative *miRNA172* binding site (Supplementary Figure 7b and c). The genomic position, expression analysis, protein sequence data, and predicted deleterious effect of the insertion in intron 8 suggest that this *APETALA2* gene is a good candidate for the *DOUBLE FLOWER* locus. *APETALA2* plays a central role in the establishment of the floral meristem and in the specification of floral organs [35, 36]. *APETALA2* was classified as a class A floral homeotic gene that specifies sepal identity if expressed by itself and specifies petal identity if expressed together with class B genes [37]. Furthermore, *AP2* supressed *AGAMOUS* expression (class C gene) in the two outer floral whorls in the floral meristem (reviewed in [38]). In rose, a reduction of *RhAGAMOUS* transcripts was proposed to be the basis of the *double flower* phenotype [39]. We can hypothesise that mis-regulation of the rose *APETALA2* homolog (due to the presence of the transposable element) might be responsible for the *RhAGAMOUS* transcript reduction, leading the *double flower* phenotype.

Interestingly, a GWAS approach using mainly tetraploid and double flower varieties [31] revealed that the most significant QTL for the number of petals (quantitative analysis) comprising most of the highly significant associated markers in a large cluster is located at the *DOUBLE FLOWER* locus (Figure 4c). Several markers in this cluster display significant dose-dependent effects on the number of petals. One of these markers, RhK5_4359_382 (at position 33.55 Mb), was analysed via the KASP technology both in the original association panel of 96 genotypes and in an independent population of 238 tetraploid varieties and showed the same effect in both populations (Supplementary Figure 8a and b). Two other markers (RhK5_14942 and RhMCRND_760_1045) were also analysed on the 96 genotypes by KASP technology and showed the same pattern (Supplementary Figure 8c and d). This demonstrates a dual role of the double flower locus for both the control of the double flower phenotype (double vs. single flowers) and the control of petal number in roses. Furthermore, the QTL for the number of petals can be detected in several association panels of unrelated rose genotypes and therefore seems to act independently of the genetic background.

#### C. Self-incompatibility

As described for other *Rosaceae* species [40–42], in some diploid roses self-incompatibility (SI) is caused by a gametophytic SI-locus. This locus is most probably composed of genes encoding S-RNAses and F-Box proteins, which represent the female and male specific components, respectively. Previous approaches have failed to characterise the rose S-locus genes due to the low sequence similarity between S-RNAse genes across species and the existence of multiple genes for both S-RNAses and F-Box proteins. A screen for S-RNAse and F-Box homologues in the HapOB genome sequence identified a region of 100 kbp on Chromosome 3 that contains three genes coding for S- RNAses and four genes for S-locus F-Box proteins (Figure 4a, Supplementary Figure 9a). This region is syntenic with the SI locus in *Prunus persica* (Supplementary Figure 9b). One of the S-RNAses (SRNase36) was expressed in pistils of ‘Old-Blush’ flowers. Of the F-Box genes, F-Box38 accumulated in stamens (Supplementary Figure 9c and 9d). Hence, this region fulfils the requirements of a functional S-locus.

This is further supported by previous data on segregation of the SI-phenotype in a diploid rose population, where the SI-phenotype had been analysed by generating a bi-parental progeny and backcrossing individual progeny to both parents [43]. We generated a marker for an orthologue of the S-RNAse gene (S-RNAse30) expressed in pistils of ‘Old Blush’ that co-segregates with the S-locus at a distance of 4.2 cM. The large number of recombinants might be explained by incomplete expression of self-incompatibility (leaky phenotypes) in some individuals of the progeny, a phenomenon that is also observed in e.g. *Solanum* populations [44].

#### D. Prickle density

We investigated the genetic determinism of prickle density in rose. In two F1 progenies, QTLs were detected on LG3. On the OW progeny, a large effect QTL was detected between position 31.2 Mb and 44.5 Mbp on male and female maps (Figure 4d). In the YW population a region was identified between 39.5 Mbp and 46.4 Mbp, which has three neighbouring QTL peak regions (Figure 4e). This region overlapped with the OW QTL (Figure 4d). Using a diversity panel, by GWAS, we were able to detect a strong association between SNPs and the presence of prickles between positions 31.0 Mbp and 32.4 Mbp (Supplementary Figure 10a). In rose, prickles originate as a deformation of glandular trichomes in combination with cells from the cortex [45]. We have looked for homologs of candidate gene controlling trichome initiation and development identified in *A. thaliana* [46]. Screening the QTL region on Chr3 of HapOW for gene family members of these candidate genes revealed several WRKY transcription factors, of which RC3G0244800 (positioned at 33.40 Mbp, Figure 4a) shows strong similarity with *AtTTG2* (*TESTA TRANSPARENT GLABRA2*), involved in trichome development in *Arabidopsis* [47] (Supplementary Figure 10b). We studied the expression of the rose *TTG2* homolog (*RcTTG2*) in three different individuals of the OW progeny with different prickle densities (absence, medium and high density prickles on the stem Supplementary Figure 10c). The *RcTTG2* transcript accumulated at higher levels in stems presenting prickles, suggesting that *RcTTG2* is a positive regulator of prickle presence in rose. This TGG2 homolog represents a good candidate for the control of prickles in rose.

## Conclusions

We have produced a high-quality reference rose genome sequence that will represent an essential resource for the rose community but also for rose breeders. Using this new reference sequence, we have analysed important structural features of the genome including centromere positions (Figure 2) and SNP and InDel frequencies (Figure 3). This reference sequence opens the way to genomics and epigenomic approaches to study important traits in response to different environments. Cultivated roses have an allopolyploid background but segregate mainly tetrasomically [10, 48]. Hence, it is a unique model for polyploidisation and Chromosome pairing mechanisms. Pentaploid dogroses have a unique meiosis [49].

Taking advantage of this new high-quality reference sequence, rose is set to become a model species to study ornamental traits. For example, rose was previously used to study scent emission, leading to the discovery of a new pathway for the synthesis of monoterpenes [50]. Here, using a combination of genomic and genetic approaches (F1 progenies and diversity panel), we have demonstrated that this new reference sequence can be used to analyse loci controlling ornamental traits: continuous flowering, double flower and prickle density (Figure 4). We have identified and characterised candidate genes for these traits. We propose that a rose *APETALA2* homologue could control the switch from simple to double flower and unexpectedly also the number of petals within double flowers. Further analyses are necessary to validate the function of these genes. The analyses were done in diploid roses but also in tetraploid roses, allowing transfer to the actual breeding materials, with a diversity panel of mostly tetraploid garden rose varieties and the tetraploid cut flower population K5. Therefore, the data can already be used for breeding by the development of diagnostic markers as we demonstrated for petal number. For this economically-crucial trait, we have developed a genetic marker that permits the prediction of petal number, which we validated on a large panel (Supplementary Figure 8). This represents a good example of how the development and release of the rose genome sequence can accelerate gains in rose breeding. A similar approach could be taken for other important traits (such as any one of the multiple scent compounds, or foliar disease resistances), leading to the development of marker assisted selection strategies in rose.

## Materials and methods

### (1) Development of haploid ‘Old Blush’ callus

Young flower buds of ‘Old Blush’ (Figure 1a) with microspores at a mid-to-late uninucleate developmental stage (Figure 1b and c) were collected in a greenhouse, wrapped in aluminium foil and stored in the dark at 4°C for 25 days. These were then surface sterilised in 70% ethanol for 30s and in sodium hypochlorite solution (2.9° active chloride) for 15 min followed by rinsing three times in double-distilled sterilised water.

Anthers were aseptically removed using binoculars, and ground in starvation B medium [51] with minor modifications (pH 6 and sorbitol 0.1 M) for 2 min using a MSE homogeniser (Measuring & Scientific Equipment Ltd., Spenser Street, London SW1E) set at 10000 rpm. Anthers were then collected on 50 μm mesh filters, covered with a fine layer of fresh modified B starvation medium and incubated for 24h at 22°C in darkness. Anthers were transferred on MS medium containing 30g L^-1^ sucrose, 0.5mg L^-1^ BAP, and 0.1mg L^-1^ NAA in 12-well culture plates. Plates were incubated in darkness at 23°C/19°C (16h/8h) taking care not to move the boxes or expose them to light during 80 days to induce somatic embryo formation. Somatic embryos were isolated from the anthers and transferred on the same medium in petri dishes with filter paper, in 4-week intervals until the production of callus (Figure 1d). Then, callus was multiplied on the same medium in the dark until enough material for DNA extraction was produced. Homozygosity was verified using ten previously described microsatellite markers [52]. Genome sizes and ploidy levels were analysed on a flow cytometer, PASIII (488 nm, 20 mW laser) (Partec, Germany). The Cystain^®^ absolute PI reagent kit (Sysmex, Germany) was used for sample preparation. *Solanum lycopersicum* ‘Stupické polni tyckove rane’ (1916 Mbp/2C) was used as an internal standard.

### (2) Genome sequencing and assembly

#### DNA extraction for PacBio and Illumina sequencing

Callus tissues of the haploid ‘Old Blush’ HapOB line was kept in the dark for 3 days prior to DNA extraction to reduce chloroplast DNA contamination. DNA extraction was performed on 1 g HapOB callus tissue as described in Daccord et al. [53]. In total approximately thirty micrograms of gDNA was obtained in several batches for preparation of three independent SMRT bell libraries. For the first library gDNA was sheared by a Megaruptor (Diagenode) device with 30 kbp settings. Sheared DNA was purified and concentrated with AMpureXP beads (Agencourt) and further used for Single Molecule Real Time (SMRTbell™) preparation according to manufacturer’s protocol (Pacific Biosciences; 20 kbp template preparation using BluePippin (Sagesscience) size selection system with 15Kb cut-off). Two additional libraries were made excluding the DNA shearing step, but with an additional initial damage repair. Size selected and isolated SMRTbell fractions were purified using AMPureXP beads and finally used for primer- and polymerase (P6) binding according to the manufacturer’s binding calculator (Pacific Biosciences). Three library DNA-Polymerase complexes were used for Magbead binding and loaded at 0.16, 0.25, and 0.20 nM on-plate concentration spending 12, 7 and 8 SMRT cells respectively. Final sequencing was done on a PacBio RS-II platform, with 345 or 360 min movie time, one cell per well protocol and C4 sequencing chemistry. Raw sequence data was imported and further processed on a SMRT Analysis Server V2.3.0.

For Illumina sequencing, approximately 200 ng gDNA was sheared in a 55μL volume using a Covaris E210 device to approximately 500 to 600 bp. One library was made using Illumina TruSeq Nano DNA Library Preparation Kit according to the manufacturer’s guidelines. Final library was quantified by Qubit fluorescence (Invitrogen) and library fragment size was assessed by Bioanalyzer High Sensitivity DNA assay (Agilent). The library was used for clustering as part of two lanes of a Paired End flowcell V4 using a Cbot device and subsequent 2*125nt Paired End sequencing on a Hiseq2500 system (Illumina). De-multiplexing of resulting data was carried out using Casava 1.8 software.

#### Genome Assembly and Polishing

All sequence data generated derived from 27 SMRT cells encompassing 19.2 Gb of reads larger than 500 bp were assembled with CANU hierarchical assembler v1.4 [15] (version release r8046). In general default settings were used except ‘corMinCoverage’, which was changed from 4 to 3, ‘minOverLapLength’ which was increased from 500 to 1000, and ‘errorRate’ adjusted to 0.015. The assembly was completed on the Dutch national e-infrastructure with the support of SURF Cooperative using 2024 cpu hours (Intel Xeon Haswell 2.6GHz) for the complete CANU process. Illumina PE (125pb) reads were mapped onto the genome assembly using BWA-MEM [54]. Pilon [55] was then used to error correct the assembly. This procedure was repeated three times iteratively.

### (3) Development of high-density genetic maps and GWAS analysis

#### Plant material

A diploid F1 population of 151 individuals (OW) was obtained by crossing *Rosa chinensis* ‘Old Blush’ (OB) and a hybrid of *R. wichurana* (Rw) obtained from Jardin de Bagatelle (Paris, France). This population was planted at the INRA Experimental Unit Horti (Beaucouzé, France).

A diploid F1 population of 174 individuals (YW) was obtained from a cross between ‘Yesterday’ x *R. wichurana* (extended population as used in [56]). This population was planted at ILVO (Melle, Belgium).

The tetraploid K5 cut rose mapping population consisted of 172 individuals obtained from a cross between P540 and P867. It was planted in Wageningen, The Netherlands and was previously used in various QTL studies [57, 58].

The association panel comprised 96 genotypes of which 87 are tetraploid, 8 triploid and one diploid and was designed in a way to reduce the genetic relatedness between genotypes [31]. Plants were cultivated in a randomised block design with three blocks comprising one clone of each genotype both in the greenhouse and at an experimental field location at Leibniz Universität Hannover, Germany. For validation of markers an independent population of 238 tetraploid varieties have been used that was cultivated in a field plot of the Federal Plant Variety Office in Hannover, Germany.

#### Genetic map construction

The construction of the different genetic maps from F1 progenies (OW, YW and K5) is described in Supplementary Methods.

#### GWAS analysis

The GWAS analyses for petal numbers and prickle density was performed in Tassel 3.0 [59] as described in Schulz et al (2016) [31]. Trait marker associations for petal number was analysed using MLM and 39831 Markers (Petal as quantitative Trait with O+K Model). Significance thresholds were corrected for multiple testing by the Bonferroni method using the number of contigs (19083) as correction factor, resulting in a significance threshold of 2.48E-6. The kinship matrix used in MLM was calculated for 10000 SNP-markers with the software SPAGeDi 1.5 (Zitat) as described in Schulz et al. 2016 [31]. For the GWAS analysis of prickles and petals with GLM in Tassel 3.0 [59], 630000 markers were analysed. Petals and prickles were set as qualitative traits (0 and 1) and analysis was performed without any correction for population (Q+K). Significance thresholds in GLM were corrected by number of contigs (28054) to 1.88E-6.

#### KASP-assay for SNP validation

SNP-markers for Kompetitive Alelle Specific PCR (KASP) assays were designed by LGC Genomics (London, UK). Genotyping was performed using a StepOnePlus Real-Time PCR system (Applied Biosystems, USA) with 20 ng DNA, 5 μl KASP V4.0 Mastermix 96/384, High Rox, 0.14 μl KASP by Design Primer Mix in a final volume of 10 μl for each reaction. KASP thermocycling was done according to the manufacturer’s standard protocol: Activation for 15 min at 94°C, followed by 10 cycles at 94°C for 20 s and 61°C for 1 min. (61°C decreasing 0.6°C per cycle to achieve final annealing temperature of 55°C) followed by 26 cycles at 94°C for 20 s and 55°C for 1 min. Reading of KASP genotyping reactions on the qPCR machine was performed in a final cycle at 30°C for 30 s. If fluorescence data did not form satisfactory clusters the conditions for additional cycles were 94°C for 20 s followed by 57°C for 1 min (up to 3 cycles). Genotypic data were analyzed with the StepOne™ Software v2.3 (Applied Biosystems, USA).

#### Development of a SCAR marker for the SI locus

The marker segregating at the SI locus was derived from a collection of genomic contigs generated by genomic sequencing of the diploid *R. multiflora* hybrid 88/124-46 via Illumina sequencing and subsequent assembly (data not shown). This collection of 27094 scaffolds was screened with 64 amino acid sequences of annotated S-RNAses from various *Prunus* species obtained from the NCBI database. One contig (contig no. 2308) with a significant hit to an S-RNAse gene also contained a predicted gene with similarity to an S-locus related F-Box protein. PCR primers (PrimerP9f 5’_CTTGCATTCAAGGTGCAGTC_3’ and Primer P9r 5’_CGGCTCTGGTGAAATAGTCC_3’) for the S-RNAse homolog were designed with the Primer 3 software and flank the intron between exon 2 and 3 of the predicted S-RNAse sequence. Amplification conditions were: 94°C for 30s, 30 cycles of 94°C for 30s, 61°C for 30s and 72°C for 2min followed by a final amplification at 72°C for 10min.

### (4) Alignment of the HapOB rose genome with the OW genetic maps

The alignment of the genetic and physical maps was done in two steps. First the HapOB sequence was aligned to the integrated genetic maps in order to detect problems of assembly (contigs present on two linkage groups). Second, to precisely order and orient the contigs on each linkage group, the alignment was done separately on the male and female maps, and manually integrated.

During the first step, 7822 out of a total of 7840 SNP markers were positioned by mapping the corresponding 70 bp probes onto the HapOB genome sequence using Blat v.35 [60]. Markers with more than one best hit were eliminated. Out of the 7360 remaining markers, 6808 passed the mapping quality filter (≥95% match, ≤4% mismatch) were retained. Of these, 6746 markers belonging to the most common linkage group on their respective contigs were conserved and described as “concordant” markers. Only contigs having more than one of these markers were retained.

During the second step, the mapping and anchoring was done independently on the male and female maps (Table 1). The procedure and conditions were the same as for the first mapping. Only concordant markers were kept (4875 (87%) and 1871(81%) for the female and male map, respectively). We positioned and oriented the different contigs manually (Supplementary Table 8). When a contig spanned several loci, its order and position was clear. However, for some contigs, genetic maps did not resolve orientation problems. In these situations, we used the synteny between *Rosa* and *Fragaria vesca* [10]. The strategy used to position and orient contigs is described in Supplementary Figure 11. The position and orientation of the contigs are listed in Supplementary Table 8.

Concerning K5 integrating genetic map, among the 25695 SNP markers present on the K5 genetic map, 20706 SNPs (80,6%) could be positioned on the HapOB genome sequence by BLAST of the SNP-flanking marker sequences (Supplementary Table 2).

### (5) Centromere region identification and fluorescent *in situ* hybridization

Three complementary tools were used to identify centromeric tandem repeats and to estimate their abundance in the *R. chinensis* ‘Old Blush’ genome: Tandem Repeat Finder (TRF, [61]), TAREAN [62] and RepeatExplorer [63], each with default settings, and the output was parsed using custom python scripts. After identification of all tandem repeats identified by TRF they were subjected to all-against-all BLAST to cluster similar repeats and to estimate abundance (total number of tandem repeat cluster copies) in the genome. Paired reads were quality filtered and trimmed to 120 bp for analysis by RepeatExplorer (0.5M read pairs) and TAREAN (1.3M read pairs). RepeatExplorer cluster CL226 had the globular-like shape specific for tandem repeats. The corresponding monomer repeat sequence was identified by analysing the contigs of this cluster with TRF. The identical tandem repeat was also identified by TAREAN and TRF. To determine the location of the CL226 tandem repeat cluster in the genome assembly, 275M paired-end genomic reads of ‘Old Blush’ were mapped onto the contigs from RepeatExplorer cluster CL226, using Bowtie2 [64] with parameter -k 1 to select read pairs with high similarity to the CL226 repeat. Selected read pairs were then split into two groups: one set of reads that match the CL226 repeat sequence itself, and the other read of that pair is placed in the group that reflects the flanking genome sequence. Both groups of reads were separately mapped onto the genomic scaffolds using Bowtie with parameters -a 1 and -N 1. The genome distribution of the two sets of CL226 reads was visualised using the circlize package [65] of R Bioconductor [66]. Mitotic Chromosome slides were prepared with the “SteamDrop” method [67] using young root meristems of *R. chinensis* ‘Old Blush’. Two oligonucleotide probes (5’-TTGCGTTGTTCTAGTGACATTCATAMRA-3’; 5’-ACCCTAGAAGCGAGAAGTTTGG-TAMRA-3’) were used for FISH, as previously described [68]. DRAWID [69] was used for Chromosome and signal analysis.

### (6) Annotation of the rose genome

Gene and TE annotations is described in Supplementary Method.

### (7) Diversity analysis

The plant material originated from ‘Loubert Nursery’ in Rosier-sur-Loire, France (*R. persica*), from ‘Rose Loubert’ rose garden in Rosier-sur-Loire, France *(R. moschata, R. xanthina spontanea* and *R. gallica)* and from ‘Roseraie du Val de Marne’, Haÿ-Les-Roses, France *(R. chinensis* var. *spontanea, R. rugosa, R. laevigata* and *R. minutifolia alba)*. Illumina paired-end shotgun indexed libraries were prepared from three μg of DNA per accession, using the TruSeq®DNA PCR-Free LT kit (Illumina). Briefly, indexed library preparation was performed with low sample protocol with a special development to reach insert size of 1-1.5 kb: DNA fragmentation was performed by AFA (Adaptive Focused Acoustics™) technology on focused-ultrasonicator E210 (Covaris), all enzymatic steps and clean up were realised according to manufacturer’s instructions, excepted fragmentation and sizing steps. According to manufacturer’s instructions, paired-end sequencing 2 × 150 sequencing by synthesis (SBS) cycles was performed on a HiSeq^®^ 2000/2500, Rapid TruSeq^®^ V2 chemistry (Illumina) running in rapid mode using on board cluster generation. For some readsets, a low enrichment of libraries with 5 cycles was performed.

Raw reads of each *Rosa* species were processed and only high quality reads were considered for further analysis. Paired end reads were mapped against the HapOB reference using BWA with default parameters [70]. Unmapped and duplicated reads were removed by SAMtools and Picard package, respectively [71]. Furthermore, reliable mapped reads were used to identify SNPs and Indels using Genome Analysis Toolkit (GATK) software [72]. To filter out the high quality SNPs, VCFtools [73] was used with minimum depth (DP) 20 and SNP quality (Q) 40. SnpEff and SnpSift [74, 75] were used to annotate the effects of SNPs and identify the potential functional effects of amino acid substitution on corresponding proteins, respectively.

To conduct, the synteny analysis between HapOB reference sequence and *Fragaria vesca*, the orthologs genes were identified using reciprocal blast with e-value 1e5, [76] v=5 and b=5. The protein sequences and annotation for *Fragaria vesca* (v2.0.a1) were downloaded from GDR database (https://www.rosaceae.org/). The output of the blast tool was used in McSCANX tool to identify syntenic regions between genomes [77]. Further, circos software [78] was used to visualise the synteny regions between two genomes. Later, the microsynteny was performed between *R. chinensis* ‘Old blush’ and *Fragaria vesca* for Chromosome 3 to see the conserved region near the RoKSN locus using Symap software [76].

Good quality and pre-processed ILLUMINA reads of *R. laevigata* were used for assembly. Genomic sequence reads were assembled using SPAdes (ver 3.11.1) at kmer value=63 [79].

### (8) Morphological traits

#### Petal number

For the OW and YW population, the number of petals per flower was counted using 5 to up to 10 independent flowers, respectively. In roses single flowers typically have 5 petals. Rose with less than 7 petals were considered as simple and with more than 8 were considered as ‘double’ flowers.

For the GWAS panel, the number of petals was counted for three flowers on each of three clones from greenhouse-grown plants and arithmetic means were calculated for each genotype.

#### Prickle number

In the OW and YW populations the length of a stem part with four internodes was measured in the middle of a stem (between 5^th^ and 7^th^ internodes). Prickles were counted on four internodes. The prickle density was expressed as the number of prickles per internode. For each genotype 3 stems were measured and counted.

For the GWAS panel, prickle density was calculated as the arithmetic mean of the number of prickles between the third and fourth node of newly-developed shoots. For each genotype, three shoots were counted from three replicates in a randomised block design.

#### Expression analysis

For *TTG2* expression analysis, three individuals of the OW progeny were selected according to prickle density: OW9068 (no prickle), OW9155 (low density) and OW9106 (high density). The terminal part of young stems were harvesting in spring 2016 from field-grown plants. RNA extraction, cDNA synthesis, qPCR and relative quantifications were performed as previously described [80]. Calibration was done using *TCTP* and *UBC* genes. The following primers were used to amplify *TTG2* (RcTTG2-1-F: CCTCAAACCCAGGAGCATC and RcTTG2-1-R: CAACAGCTTGATCCCTGAGAG). Organ-specific expression of candidate self-incompatibility genes were tested using RNAs derived from stamens and pistils of 3 flower buds and 5 open flowers, and terminal leaflets of 3 young leaves, sampled from an individual of ‘Old Blush’ in August 2017. RNA extraction was carried out according to previous protocols [39]. cDNA synthesis and RT-PCR were performed with PrimeScript RT reagent Kit with gDNA Eraser and EmeraldAmp PCR Master Mix (TaKaRa^®^, Japan) according to the manufacturer’s protocols. The following primers (5’ to 3’) were used to amplify 7 candidate genes and a house-keeping gene: *S-RNase 26* (F1: TGCAGCCAACACATACGATT and R1: GCAAGAAGATCGGCGTAGTC), *S-RNase 30* (F1: TGTTCAACAATGGCCGATAA and R1: TGCACATAAGCGAAGGAGTG), *S-RNase 36* (F1: TGTGGTAACAGCTGCAAAGC and R1: TCAACCACGTTTTTGCCATA), *F-box 29* (F2: TGACTATTTTCTATTGCGCTTGAG and R1: CACCACAAAAAGGATAACAAGAC), *F-box 31* (F1: TTTGCTATGAAAATGATAACAACAG and R1: AACCCCATGGTTTCATTAAGTA), *F-box 38* (F1: GACTACTCTCCTTTGGCCTGAA and R1: CTACAGCTGCAGAATCATTTGAC), *F-box 40* (F1: CGTCCAATATCTCTACTCAATGGT and R1: CCTCTTCTTGGTGAGTCTGAAAT), *RoTCTP* (F2: AAGAAGCAGTTTGTCACATGG and R2: TCTTAGCACTTGACCTCCTTCA).

## List of abbreviations

Transposable Elements (TE); Fluorescent *in situ* Hybridization (FISH); Single Nucleotide Polymorphism (SNP); 4’,6-diamidino-2-phenlylindole (DAPI); TESTA TEGUMENTA GLABROUS2 (TTG2), Genotyping By Sequencing (GBS); Kompetitive Allele Specific PCR (KASP); Once-Flowering (OF), Continuous-Flowering (CF), Genome Wide Association Study (GWAS); quantitative Polymerase Chain Reaction (qPCR); Reverse-Transcriptase Polymerase Chain Reaction (RT-PCR); SSR: Simple Sequenced Repeat; QTL: Quantitative Trait Locus

## Additional files

### Table

**Table 1:** Table 1.docx (word)

Metrics of the alignment of the male and female genetic maps with the HapOB genome assembly. The genetic maps were developed from a cross between ‘Old Blush’ (female) and a hybrid of R. wichurana (male) using a Affymetrix SNP array. The initial size of the genome was 512Mb, and reached finally 518,5 due to the addition of 10000N between each contigs to create the pseudo-molecules.

**Table 2:** Table2.docx (word)

Summary of resequencing and variations (SNPs and Small Indels) indentifed in 8 *Rosa* species

### Supplementary Figures

**Supplementary Figure 1**

Alignment of the OW and YW genetic maps to the pseudo-chromosomes of the HapOB physical sequence. Per mapping population, female (black circles) and male (gray circles) genetic maps are shown separately. Vertical bars show the location of OBC226 repeat sequence (red) and pericentromeric region (gray).

**Supplementary Figure 2a**

Alignment of the physical sequence of the seven pseudo-chromosomes of HapOB to the K5 integrated genetic map [70]. Several contigs that are currently assigned to Chr0 can be anchored to the different K5 linkage groups. LG3 of the K5 genetic map was inverted to be in the same orientation as the physical sequence of Chr3 of HapOB.

**Supplementary Figure 2b**

Alignment of the physical sequence of the seven pseudo-chromosomes of HapOB to the YW integrated genetic map. Several contigs that are currently assigned to Chr0 can be anchored to the different YW linkage groups.

**Supplementary Figure 3**

Abundance of repetitive elements in the HapOB genome. The value represents the cumulative genome coverage of the TE family in percentage of the total genome size. The numbers represents the number of copies for each transposable element family of the consensus library. Details are presented in Supplementary Table 3.

**Supplementary Figure 4**

Synteny analysis between the HapOB genome and the woodland strawberry genome *(Fragaria vesca*). Rh and Fv for the seven rose and strawberry chromosomes respectively.

**Supplementary Figure 5**

Analysis of the *RoKSN^null^* allele in HapOB. The upper part of the figure represents the macrosynteny analysis between the *CONTINUOUS FLOWERING* locus in HapOB and *Fragaria vesca*. Large segmental rearrangements are detected between conserved blocks (A, B, C and D) and no *RoKSN* homolog is present in HapOB. The lower part represents the microsynteny at the *FvKSN* locus between *F. vesca* and once-flowering roses (*R. multiflora* and R. *laevigata*). In these once-flowering roses, the synteny is conserved (no rearrangement) and the *RoKSN* gene is present (shaded area in lower panel). The *KSN* homologues are shown in red (gene30276 is *FvKSN*). Other orthologous genes are connected by black lines.

**Supplementary Figure 6**

Amino-acid alignment of the protein encoded by candidate-gene at the *DOUBLE-FLOWER* locus. A) Amino-acid alignment of the F-Box protein encoded by the two alleles of ‘OldBlush’ gene RC3G0245100. 1. HapOB: the allele present in the HapOB reference sequence; 2. OB2: the second allele of ‘OldBlush’. B) Amino-acid alignment of the tetratricopepetide repeat (TPR)-like family protein encoded by the two alleles of ‘OldBlush’ gene RC3G0243500. 1. HapOB: the allele present in the HapOB reference sequence; 2. OB2: the second allele of ‘OldBlush’. C) Amino-acid alignment of the protein with high similarity to the Ypt/Rab-GAP domain of the gyp1p super family encoded by the two alleles of ‘OldBlush’ gene RC3G0245000. 1. HapOB: the allele present in the HapOB reference sequence; 2. OB2: the second allele of ‘OldBlush’.

**Supplementary Figure 7**

Analysis of the two *APETALA2* alleles in ‘Old Blush’. A) Genomic organisation of the two alleles. The gene contains 10 exons (numbered yellox boxes show CDS) and 9 introns. The main difference between both alleles is the insertion of a large element of unknown length in intron 8 of one Old Blush *AP2* allele, which are flanked by regions with homology to repetitive elements (TE). B) Conservation of two AP2 domains in APETALA2 of *Arabidopsis thaliana* (*At*) and rose (Rc). C) Conservation of the *miRNA172* binding sequences in the *Arabidopsis* APETALA2 clade genes (*AP2, TOE1, TOE2* and *TOE3*) and the two alleles of the *APETALA2* homolog in rose. D) Phylogenetic analysis of the AP2 and ANT clades of the APETALA2 protein subfamily of rose and A. *thaliana*. The *Arabidopsis* proteins are numbered according to TAIR nomenclature (http://www.arabidopsis.org).

**Supplementary Figure 8**

Validation of SNP markers for petal number. A-B) Marker RhK5_4359_382 (at position 33.55 Mbp) in an association panel of 96 cultivars (A), and in 238 independent tetraploid rose cultivars (B). C) Marker RhK5_14942 (at Chr3 position 33.24 Mbp) in an association panel of 96 cultivars. D) Marker RhMCRND_760_1045 (at Chr3 position 33.21 Mbp) in an association panel of 96 cultivars. Numbers on x-axis show allele dosage for the four marker classes (0 and 4 for the alternative homozygotes, and 1-3 for the heterozygotes). Per allele dosage group, the number of individuals (n) is given on top; groups that are significantly different at p ≤ 0.05 are indicated by letters above the whiskers. The mean is represented by small white squares, and the median by the horizontal line. Mean values are given above the box. Whiskers represent the standard deviation and box size the standard error.

**Supplementary Figure 9**

Candidate genomic region of the self-incompatibility locus on Chromosome 3 (Chr3) of HapOB. A) Location of candidate genes in a 100 kbp region of the rose S-locus; *S-RNase* and *F-Box* genes are depicted as black arrows, other genes as gray arrows. B) synteny between the genomic regions surrounding the S-locus in peach (red diamond, Chr6) and rose (Chr3). C) Tissues from flower bud and open flower sampled for expression analysis by RT-PCR. D) RT-PCR analyses of candidate genes (*S-RNase* and *F-Box*). St for stamen; Pi for pistils; L for leaves. Genomic DNA (gDNA) is used as positive control, and *RoTCTP* is used as house-keeping gene.

**Supplementary Figure 10**

A *TTG2* homolog is a candidate gene for the control of prickle. A) GWAS analysis showing p-values of the association between SNPs positioned along Chr3 and the absence or presence of prickles. Significant SNPs are located between positions 31 Mbp and 32.4 Mbp. B) Phylogenetic analysis of the Arabidopsis *TTG2* clade of the *WRKY* transcription factor family. The rose WRKY transcription factors located in the prickle density QTL region are shown in green and the closest *TTG2* homolog is shown in red. The *Arabidopsis* WRKY proteins are numbered according to TAIR nomenclature (http://www.arabidopsis.org). C) *TTG2* transcript accumulation in three different OW individuals with no prickles, medium, and high prickle density. The transcript accumulation level was analysed by qPCR and expressed as a ratio relative to the sample without prickle.

**Supplementary Figure 11**

Strategy used to position and orient the contigs by anchoring onto the OW genetic map, illustrated on the upper half of LG5, for the male map (top) and for the female map (bottom). Genetic loci used for anchoring are indicated by gray tick marks (on average 8 SNP markers per locus). Per contig (boxes), sequence orientation is indicated by the orientation of the contig number within the box. Contigs that are only anchored to one locus (e.g. contig 528, 561, 2014 or 2101), are oriented based on synteny with *Fragaria vesca*. Dashed lines connect contigs anchored to the female and male maps. The final order of the contigs is presented with the contigs orientation. During anchoring, the following cases were encountered. Case 1: The contig is not oriented with the male map, but can be oriented thanks to the female map (e.g. contig 2014, orientation -). Case 2: The contig is anchored only on the male or female map (e.g. contig 375). We manually integrated the contig upstream of contig 2085. Case 3: At a same locus, more than one contig is anchored (e.g. contig 561 and 2101 on the female map). In order to position and orient the contigs, we used synteny with *Fragaria vesca*, and positioned contig 2101 (orientation -) before contig 561 (orientation +), and both between contigs 2099 and 2012

### Supplementary Tables

**Supplementary Table 1**

Validation of the homozygocity of HapOB using ten microsatellites located on the 7 linkage groups

**Supplementary Table 2**

Details of the high density SNP genetic maps from the OW progeny, obtained from a cross between *R. chinensis* ‘Old Blush’ (OB, female) and a hybrid of *R. wichurana* (W, male). Linkage maps for the maternal (A1 to A7) and paternal (B1 to B7) parents are provided, specifying the number of SNP per LG, the size (in cM), the number of unique loci per LG and the density of SNP/cM.

**Supplementary Table 3:** SupplementaryTable3.xlsx (Excel file)

Consensus librairy for Transposable Elements annotation in HapOB genome. Each line reprensents a TE consensus genome family. TE consensus name: name of the consensus. Length (bp): TE consensus length in bp. Coverage (bp): cumulative genome coverage of this TE family in bp. Coverage (%): cumulative genome coverage of this TE family in percentage (calculated on 518515953 bp). frags: number of TE fragments before the TEannot “long join procedure”. fullLgthFrags: number of complete TE fragments before the TEannot “long join procedure” (a full-length fragment represent only one fragment that covers 95% of the consensus), copies: number of TE copies after the TEannot “long join procedure” (a copy is a chain of fragments). fullLgthCopies: number of complete TE copies after the TEannot “long join procedure” (a full-length copy is reconstructed by the join of fragmented multiple hits and all these fragments cover 95% of the consensus), meanld: mean identity calculated on the percentages of identity (between each copy with its TE consensus), sdId: standard deviation on the percentages of identities.

**Supplementary Table 4**

SNP analysis in resequenced species within the genus *Rosa*. For each species, the detected SNP are classified according to their effect (HIGH (non synonymous change, splice sites), LOW (synonymous), MODERATE and MODIFIER (intron, intergenic)) or to their position in the genome (coding gene, intergenic).

**Supplementary Table 5**

Detection of the *RoKSN^null^* allele in *R. chinensis* ‘Old Blush’ using two different progenies. We studied the *RoKSN* segregation in two different progenies: (a) *R. chinensis* ‘Old Blush’ X a hybrid of *R. wichurana* (*RoKSN^WT^/RoKSN^copia^*, OW progeny) and (b) *R. chinensis* ‘Old Blush’ X *R. moschata (RoKSN^WT^/RoKSN^WT^*, OM progeny). In both populations we observed unexpected allelic combinations, with individuals presenting only the *RoKSN^WT^* allele (47 individuals out of 152 in OW and 5 individuals out of 10 in OM). One hypothesis to explain these unexpected results is the existence of a null allele in *R. chinensis* ‘Old Blush’. In other words, our data suggests ‘Old Blush’ is not *RoKSN^copia^/RoKSN^copia^*, but *RoKSN^copia^/RoKSN^null^*.

**Supplementary Table 6**

Precise location of the *DOUBLE FLOWER* locus using OW progenies with 151 individuals (OW151, in orange), with 260 individuals (OW260, in yellow), the HW (H190 x hybrid of *R. wichurana*, in purple), the 94/1 (in grey) and the YW (in blue) F1 progenies.

**Supplementary Table 7**

Expression pattern during the floral development of the 41 candidate genes located in the *DOUBLE FLOWER* locus interval (see Supplementary Table 6). Expression data from Dubois et al. (2012). The expression value is expressed as the count number from RNASeq data obtained from Dubois et al. (2012): IFL for Floral Bud and Floral Meristem transition; IMO for Floral Meristem and Early Floral organs (Sepal, petal, stamens and carpel) developments; BFL for closed flower and OFT for open flower. The four most interesting candidate-genes according to their expression patterns are in blue and bold (see details in the text).

**Supplementary Table 8:** SupplementaryTable8.xlsx (Excel file)

Positionning and ordering of the contigs on the 7 pseudo-molecules. 196 contigs were anchored to the female and male genetic maps (more than one marker). Procedure for the positionning and ordering is explained in Materials and Methods, and an example (Linkage group 5) is presented in Supplementary Figure 11. The average genetic position is presented for the male and female maps in cM. The contigs in red are those for which a manual analysis as been done as described in Materials and Methods.

## Acknowledgements

We thank the ImHorPhen team of IRHS and the experimental unit (UE Horti) for their technical assistance in plant management. We thank the PTM ANAN (Muriel Bahut) of the SFR Quasav and the Gentyane platforms (especially Charles Poncet) for the SSR and SNPs analyses respectively. We acknowledge Aurelie Chauveau and Isabelle Le Clainche for libraries preparation and Elodie Marquand and Aurélie Canaguier for data processing. This work was supported by CEA-IG/CNG, by conducting the DNA QC and by providing access to INRA-EPGV group for their Illumina Sequencing Platform. We acknowledge Jean-Luc Gaignard (from the communication service of INRA) for his help to fund the project.

We thank ‘Région Pays de la Loire’ for funding the sequencing of HapOB (Rose genome project), the resequencing of eight wild species (Genorose project in the framework of RFI Objectif Végétal’) and for the EPICENTER ConnecTalent grant of the Pays de la Loire (N.D. and E.B.). F.F. and L.H.S.O. thank ANR for funding the genetic determinism of flower development (ANR-13-BSV7-0014), K.K. thanks JSPS for funding the analysis of S-locus (JSPS KAKENHI No.17H04616). T.D. thanks the German ministry of economic affairs for funding GWAS analysis (Aif programme ZI) and the Deutsche Forschungsgemeinschaft for the RNA-Seq data generation (DFG program GRK1798). The development of the high-density SNP maps was partly funded by TTI Green Genetics and by the TKI project “A genetic analysis pipeline for polyploid crops” (BO-26.03-002-001).

## Materials & correspondence

Any request for correspondence and materials should be sent to Fabrice Foucher.

## Data availability

All the genome data have been made available on a genome browser (https://iris.angers.inra.fr/obh/). Fasta files of chromosomes and genes (mRNA, Proteins and ncRNA) and gff files for gene models and structural features (TE) can be downloaded. RNASeq data used for genome annotation are available under the following SRA accession (SRP128461 for 91/100-5 leaves infected with balckspot and for *R. wichurana* and Yesterday leaves infected with two powdery mildew pathotypes). Raw data of resequencing of the eight wild *Rosa* species are available under the SRA accession number SUB3466405

## Author contributions

LHSO developed the OW genetic map, analyzed the haploid, performed genetic determinism studies on the OW progeny. IK performed and interpreted the analyses of centromeric regions. KVL performed FISH analysis. LL performed cytometric analysis of the HapOB line. TR, LL and JDR developed the YW genetic map. JDR and TR aligned the YW genetic map to the HapOB reference sequence. JDR and LL performed QTL analyses on prickles and flower traits in YW and TR analyzed candidate genes in QTLs. LH developed the haploid line. LD performed the synteny and diversity analyses. PMB developed and aligned the K5 map to the HapOB reference sequence and analyzed Chr0. ZNN analyzed the genetic basis of prickle density and studied the *TTG2* candidate gene. ND performed sequence polishing and anchoring of the reference sequence to the OW genetic map. DS, NE and ML contributed to the GWAS approach and developed KASP markers. EN generated part of the RNA-Seq data. SB produced haploid DNA for sequencing. TT developed and maintained the F1 OW individuals. AC analyzed the SNP data of the OW progeny. JJ analyzed candidate genes for double flower. LV contributed to the production of the haploid. SG developed the genome browser. TJAB and PA contributed to the development of the K5 genetic map and its alignment to the reference sequence. REV and CM contributed to the K5 and OW genetic maps. HGV, TH and ES performed rose genome sequencing and assembly. MCLP, AB and RB performed wild species re-sequencing. JC coordinated diversity analysis. NC and HQ performed the TE annotation. SA performed the gene annotation. KK performed the SI locus analysis. SS contributed to financial support and discussion for the haploid line development. MJMS contributed to the K5 analysis and to the management of the project. TD developed the GWAS approach and some of the RNA-Seq experiments, contributed to the genetic determinism analysis (double flower and SI locus) and to the management of the project. EB managed the haploid sequencing. FF performed AP2 analysis and genome anchoring to the OW genetic map, coordinated the project and the writing of the manuscript. FF, LHSO, TR, PMB, MJMS, TD and JDR were major contributors to the writing of the manuscript. All authors read and approved the final manuscript.

